# The challenge of chromatin model comparison and validation - a project from the first international 4D Nucleome Hackathon

**DOI:** 10.1101/2024.10.02.616241

**Authors:** Jędrzej Kubica, Sevastianos Korsak, Ariana Brenner Clerkin, David Kouřil, Dvir Schirman, Anurupa Devi Yadavalli, Krzysztof Banecki, Michał Kadlof, Ben Busby, Dariusz Plewczynski

## Abstract

The computational modeling of chromatin structure is highly complex and challenging due to the hierarchical organization of chromatin, which reflects its diverse biophysical principles, as well as inherent dynamism, which underlies its complexity. The variety of methods for chromatin structure modeling, which are based on different approaches, assumptions and scales of modeling, suggests that there is a necessity for a comprehensive benchmark. This inspired us to conduct a project at the NIH-funded 4D Nucleome Hackathon on March 18-21, 2024 at The University of Washington in Seattle, USA. The hackathon provided an amazing opportunity to gather an international, multi-institutional and unbiased group of experts to discuss, understand and undertake the challenges of chromatin model comparison and validation. These challenges seem straightforward in theory, however in practice, they are challenging and ambiguous. To address them, we developed a bioinformatics workflow for chromatin model comparison and validation, in which we use distance matrices to represent chromatin models, and we calculate Spearman correlation coefficients between pairs of matrices to estimate correlations between models, as well as between models and experimental data. During the 4-day hackathon, we tested our workflow on several distinct software packages for chromatin structure modeling and we discovered several challenges that include: 1) different aspects of chromatin biophysics and scales complicate model comparisons, 2) expertise in biology, bioinformatics, and physics is necessary to conduct a comprehensive research on chromatin structure, 3) bioinformatic software, which is often developed in academic settings, is characterized by insufficient support and documentation. Therefore, our work constitutes a way to advance the modeling of the 3D organization of the human genome, while emphasizing the importance of establishing guidelines for software development and standardization.

## Introduction

### Hierarchical organization of chromatin structure

Modeling 3D chromatin structure requires an examination of its multi-scale organization. At the fundamental level, DNA wraps approximately 1.65 times around an octamer of histone proteins to form nucleosomes. (Luger et al., 1997; Zhang and Huang 2021). These nucleosomes play pivotal roles in templating many biophysical processes through histone modifications (Scharf 2009). Despite previous conjectures regarding the formation of a 30 nm fiber through nucleosome clustering, contemporary consensus does not support this 30 nm modeling *in vivo* (Scharf 2009; Ricci et al., 2015; Ou et al., 2017; Wako et al. 2020; Xu et al. 2021; Zhurkin and Norouzi 2021). The intermediate scale encompasses loops and topologically associated domains (TADs), predominantly governed by two principal proteins: structural maintenance of chromosomes (SMC) complexes (Agarwal et al. 2023), characterized by ring-like configurations facilitating loop extrusion within chromatin, and CCCTC-binding factor (CTCF) loops (Lazniewski et al. 2019), exhibiting prolonged lifespans while binding to specific motifs. Consequently, SMC complexes assume a ring-like conformation to extrude loops, while CTCF functions as orientation-dependent impediments to the former. At the sub-megabase level, the communication between these loops is restricted by the topologically associated domains enabling cell-type specific gene expression programs. At the compartment level, chromatin segregates into two compartments: open (A) which is loosely arranged and biophysically accessible to transcription factors, and closed (B) which is denser (Odenheimer, Kreth, and Heermann 2005). Typically, compartmentalization is modeled employing long-range non-bonding forces, such as block-copolymer potentials (Zhou and Gao 2020), representing an amalgamation of diverse smaller-scale interactions. Finally, the human genomic landscape is demarcated into 23 pairs of chromosomes, each resembling an independent polymer chain (Kloc and Kubiak 2022). These chromosomes are densely packed in the nucleus, with specific regions (lamina-associated domains) exhibiting proximity to the nuclear lamina (Kloc and Kubiak 2022; Tolokh et al. 2023).

Consequently, it appears that each scale of the chromatin organization functions as an autonomous biological apparatus. Intriguingly, despite this autonomy, inter-scale interactions have been observed (Bártová et al. 2008; Rao et al. 2014). For instance, histone modifications are purportedly instrumental in shaping regions of open and closed chromatin, while compartmentalization correlates with lamina domains, evidenced by the propensity of B compartments to interact with the lamina (Attar et al. 2024). In conclusion, the intricate interplay of diverse proteins dynamically interacting with chromatin, combined with the complexity at each level of organization, underscores the multifaceted nature of chromatin structure.

With a rudimentary comprehension of chromatin structure, two observations emerge: its hierarchical organization manifests distinct biophysical principles, and its dynamism underscores its complexity. Addressing the former, various experimental methodologies have been devised to probe each scale individually. For instance, MNase-seq and ATAC-seq data (Chereji, Bryson, and Henikoff 2019; Grandi et al. 2022) determine the nucleosome positioning. ChIA-PET and Hi-ChIP experiments (Li et al. 2010; Yan et al. 2014) produce contact matrices and prove efficacious in TAD-scale modeling, while Hi-C experiments (Rao et al. 2014) identify compartments and subcompartments, thus an integration thereof can be useful in chromatin modeling. Nonetheless, a critical consideration often overlooked is the population and cell-cycle averaging inherent in many of these datasets, necessitating the adoption of single-cell experimental techniques such as single-cell Hi-C to mitigate this limitation (Nagano et al., 2013). Although many experimental methods were developed, there remains a lack of sufficient data that hinders a full understanding of chromatin structure. This data gap presents a new challenge: comparison and validation of 3D modeling techniques.

### Modeling of the chromatin structure

Theoretical modeling of the chromatin structure is highly complex. It requires consideration of various factors that influence the final configuration of the polymer, as well as its multi-scale organization. At the lowest level, short-range interactions between residue pairs are predominant. However, at higher levels, weaker long-range interactions maintain the compactness of the polymer in the nucleus. Therefore, incorporating all available information is essential for generating realistic models. Furthermore, computational resources and time constitute a significant challenge - more input data necessitates more computational power, which is currently limited. In addition, chromatin modeling faces other obstacles, such as a lack of method standardization and evaluation metrics, proper model visualization, and dealing with experimental data averaged over time and population. To address those challenges various approaches have been developed in recent years to model chromatin structure at different scales and resolutions. However, despite these efforts, a gold standard is still lacking, primarily because model validation against experimental data remains difficult.

### Methods for chromatin structure modeling

Strategies for chromatin structure modeling can be divided into data-driven and predictive (Fig. 2). The data-driven strategies take as input either experimental genomic data (e.g., Hi-C or ChIA-PET that provide contact frequencies) or imaging data (e.g., FISH that shows polymer density) which serves as a starting point for chromatin polymer modeling. Conversely, predictive strategies, propelled by advancements in deep learning, analyze linear data from ChIP-seq, ATAC-seq, or DNA sequences, encompassing epigenetic modifications or chromatin accessibility, to infer chromatin structure (Valeyre et al., 2022; Polovnikov et al., 2023). Output may include a contact map, a 3D model or an ensemble of 3D models, categorized by the scale of modeling, such as loops, TADs, or the whole genome. Input data can be derived from “bulk” or single-cell experiments, necessitating different considerations for model expectations. Constructing models from “bulk” Hi-C data presents challenges due to averaging chromatin states, overlooking the intrinsic heterogeneity of the underlying chromatin conformation changes. To alleviate those problems a slightly different set of methods was developed that model the chromatin conformation in particular cell-based states based on single-cell Hi-C (scHi-C) data first introduced by Nagano et al., 2013. Those methods were meant to deal with the specific focus on the sparsity of the input data which is the main problem of scHi-C. Therefore they apply methodologies highly robust to the data sparsity. Several such methods have already been implemented (Paulsen et al., 2015, Hiram et al., 2016, Carstens et al., 2016, Stevens et al., 2017, Zhu et al., 2019, Rosenthal et al., 2019, Zha et al., 2021, Meng et al., 2021). They incorporate a set of very different approaches to the problem. Most of the methods use scoring functions which are then optimized by simulated annealing protocols or gradient descent optimization (Rosenthal et al. 2019, Meng et al., 2021). Other methods define the posterior Bayesian probability function and apply Markov Chain Monte Carlo (MCMC) algorithms to draw models from the distribution (Zhu et al., 2019, Zha et al., 2021) or opt for molecular dynamics simulations (Stevens et al., 2017). Most of those methods are written either in C++ or Python and are easily accessible and usable, which makes them helpful in studying chromatin conformations at a single-cell resolution.

**Fig. 1.**
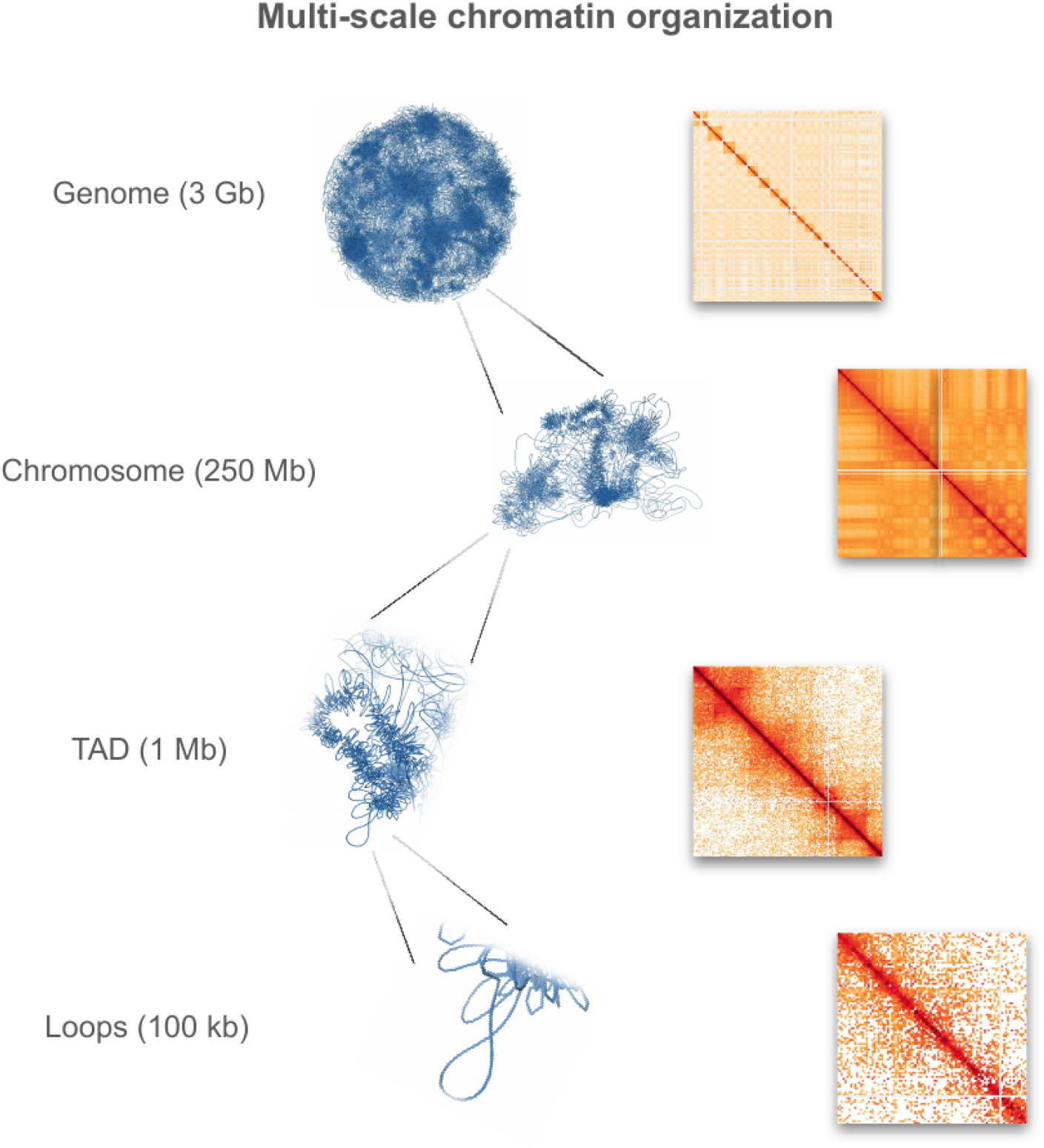
An illustration of the multi-scale chromatin organization. The approximate genome lengths (in base pairs) are presented alongside structural models and experimental contact maps for each scale (data: 4DN Data Portal *4DNES4AABNEZ* - *in situ* Hi-C on human embryonic stem cells (H1) treated with RNase A). Abbreviation: TAD - topologically associated domain.

**Fig. 2.**
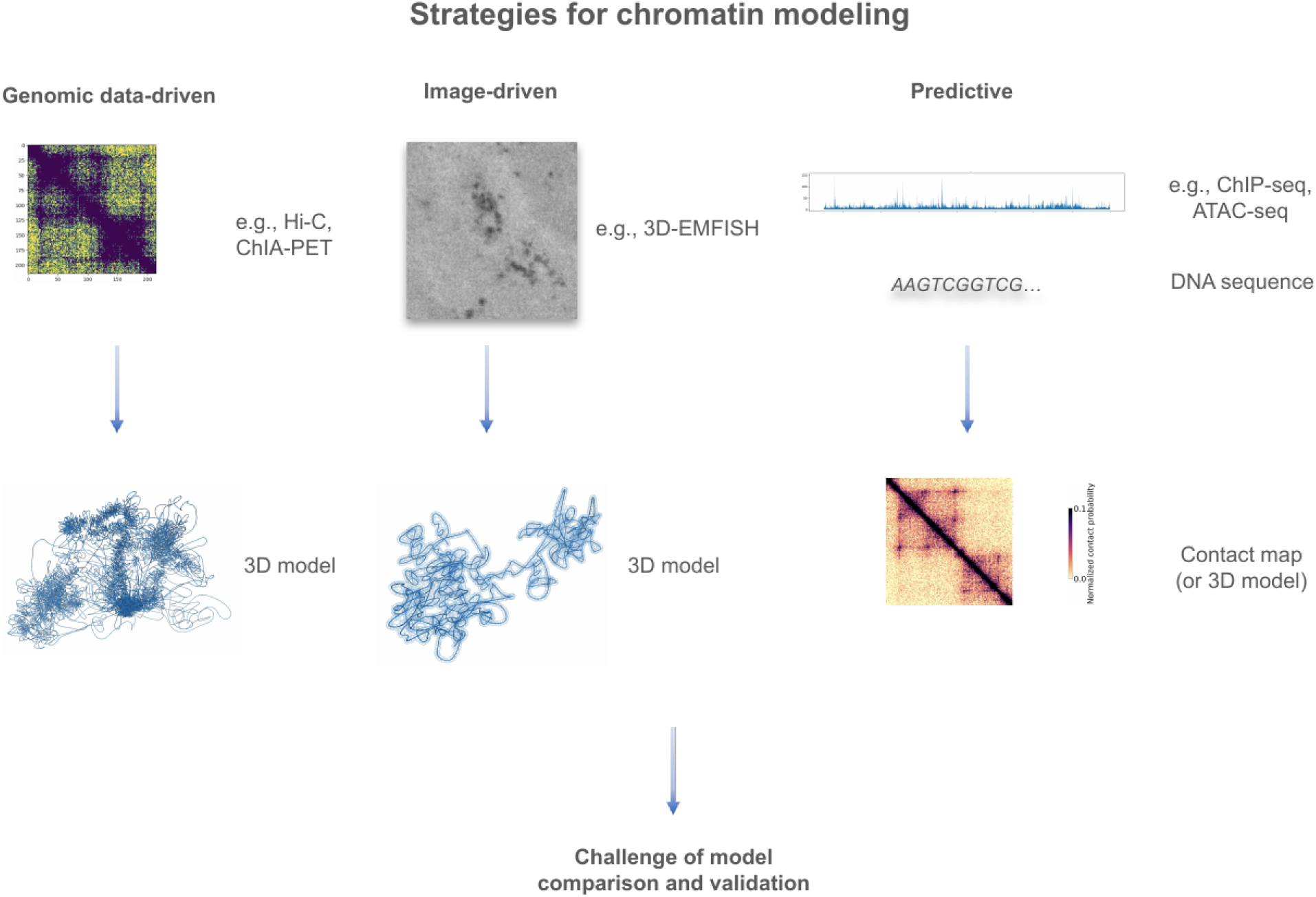
An overview of the strategies for chromatin structure modeling. Strategies for chromatin modeling can be divided into genomic data-driven, image-driven and predictive. Each strategy can produce a structural 3D model or a contact map.

### Bottom-up versus top-down modeling

Primarily, bottom-up approaches employ the first principle assumptions regarding the system’s force field to reconstruct the chromatin conformation (e.g., Spring model, MultiMM) (Kadlof, Rozycka, and Plewczynski 2020; Korsak, Banecki and Plewczynski 2024). These models incorporate loops derived from Hi-C or ChIA-PET data, utilizing virtual springs to bring spatially distant chromatin regions into proximity. Conversely, top-down models (e.g., MiChroM, GEM-reconstruction, PHi-C, miniMDS) (Di Pierro et al. 2017; Le Treut, Képès, and Orland 2018; Shinkai et al. 2020; Rieber and Mahony 2017) prioritize the optimization of model hyperparameters to emulate experimental Hi-C heatmaps, ensuring strong correlation between all-versus-all distances of the polymer and Hi-C data, rather than emphasizing the force field. Moreover, stochastic models (e.g., MoDLE, LoopSage) (Rossini et al. 2022; Korsak and Plewczynski 2024) generate thermodynamic ensembles of models, wherein the average all-versus-all distances replicate TAD structures. These models, characterized by escalating complexity, endeavor to reconstruct heatmaps by incorporating biophysical assumptions regarding loop extrusion and modeling dynamic trajectories over time. Minimalistic data, such as anchors from Hi-C, ChIA-PET, or Hi-ChIP experiments, or information gleaned from ChIP-seq experiments, are utilized to infer the locations and orientations of barrier CTCF proteins. Therefore, successful modeling necessitates the adjustment of numerous biophysical parameters about loop extrusion dynamics. Below, we present a general overview of the variety of approaches in the software for chromatin structure modeling (Table 1), which were also described in detail in recent literature on this topic (Oluwadare et al., 2019; Belokopytova & Fishman, 2021; Tao et al., 2021; Zhang et al., 2024).

**Table 1.**
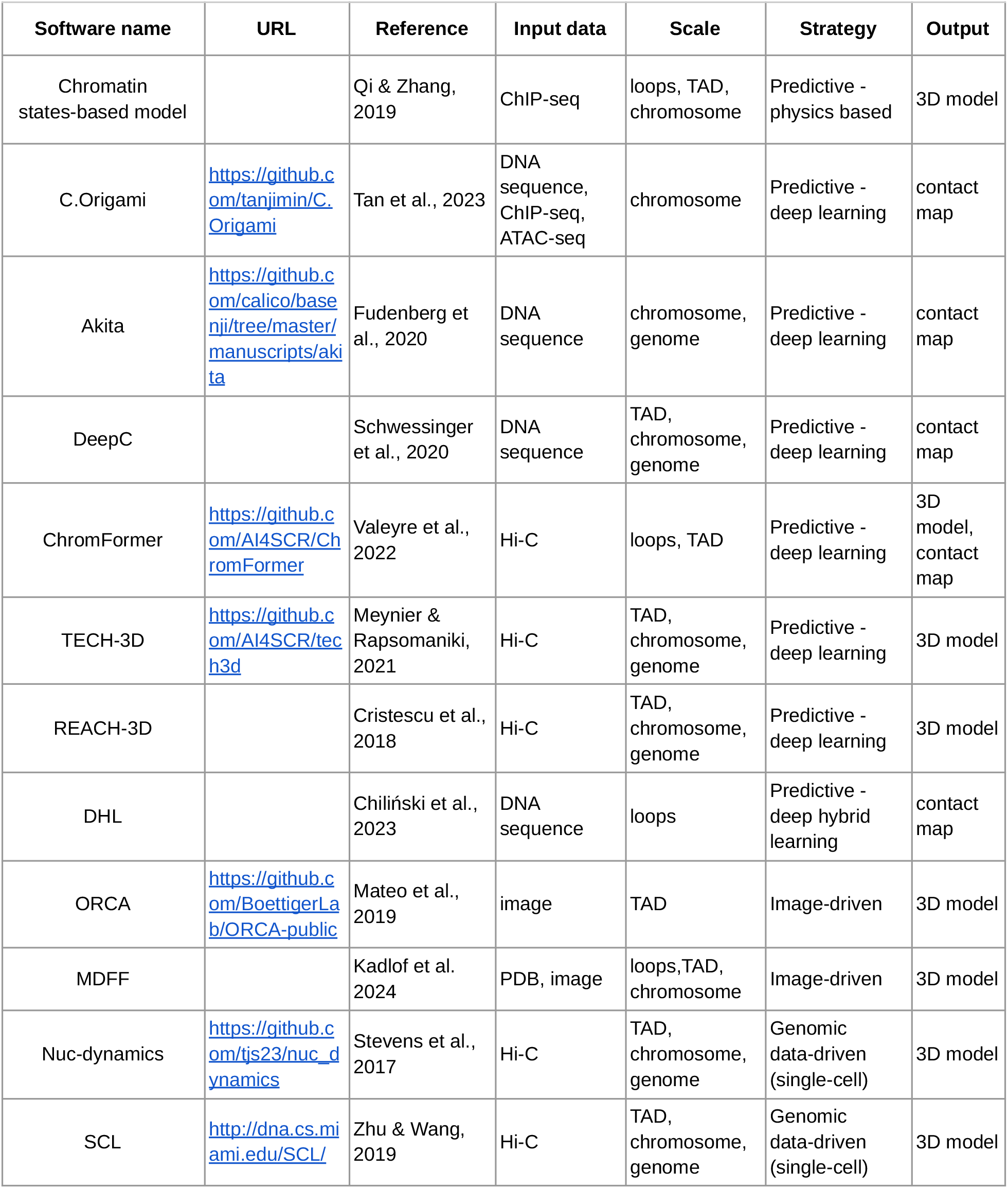
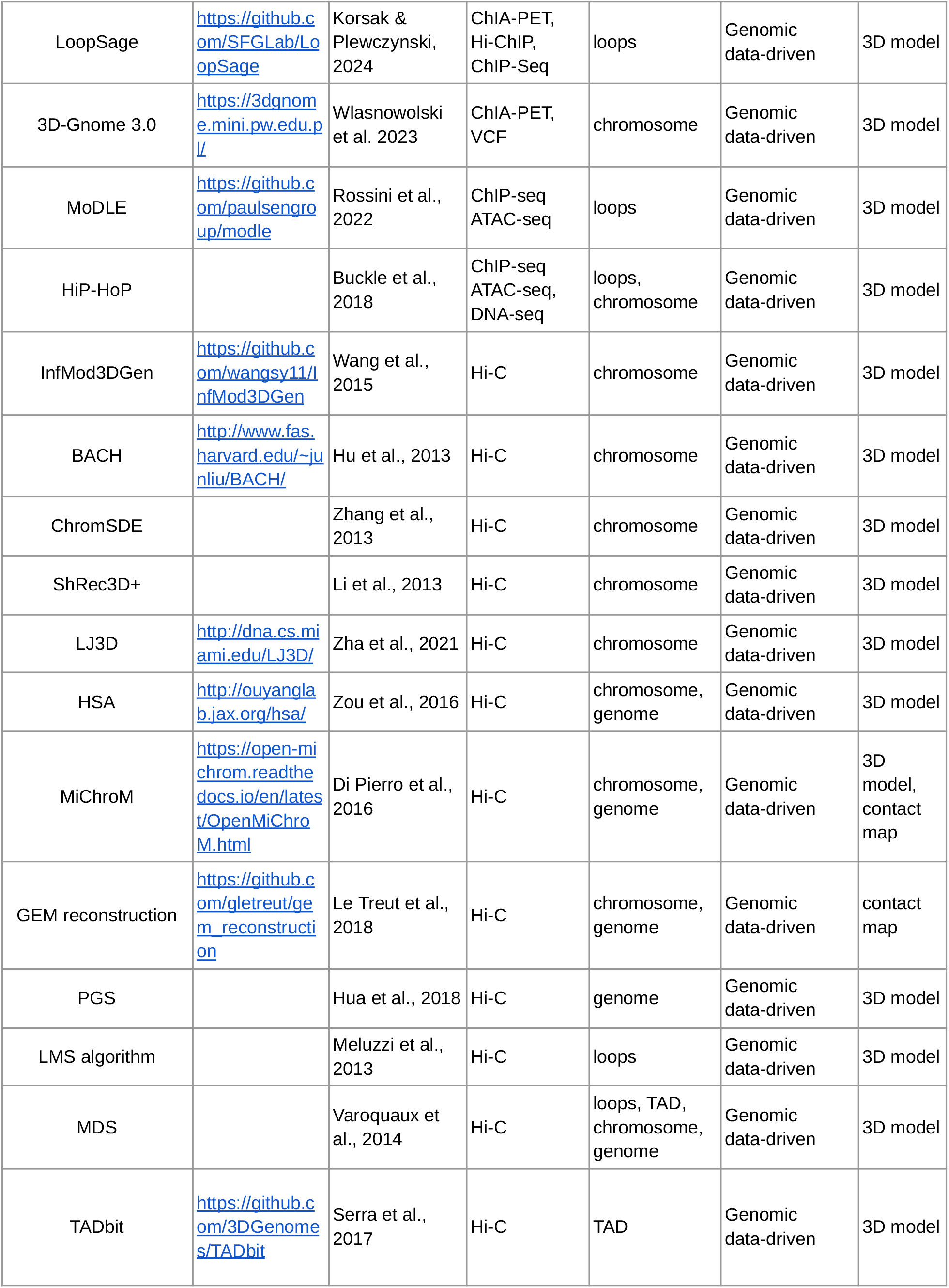
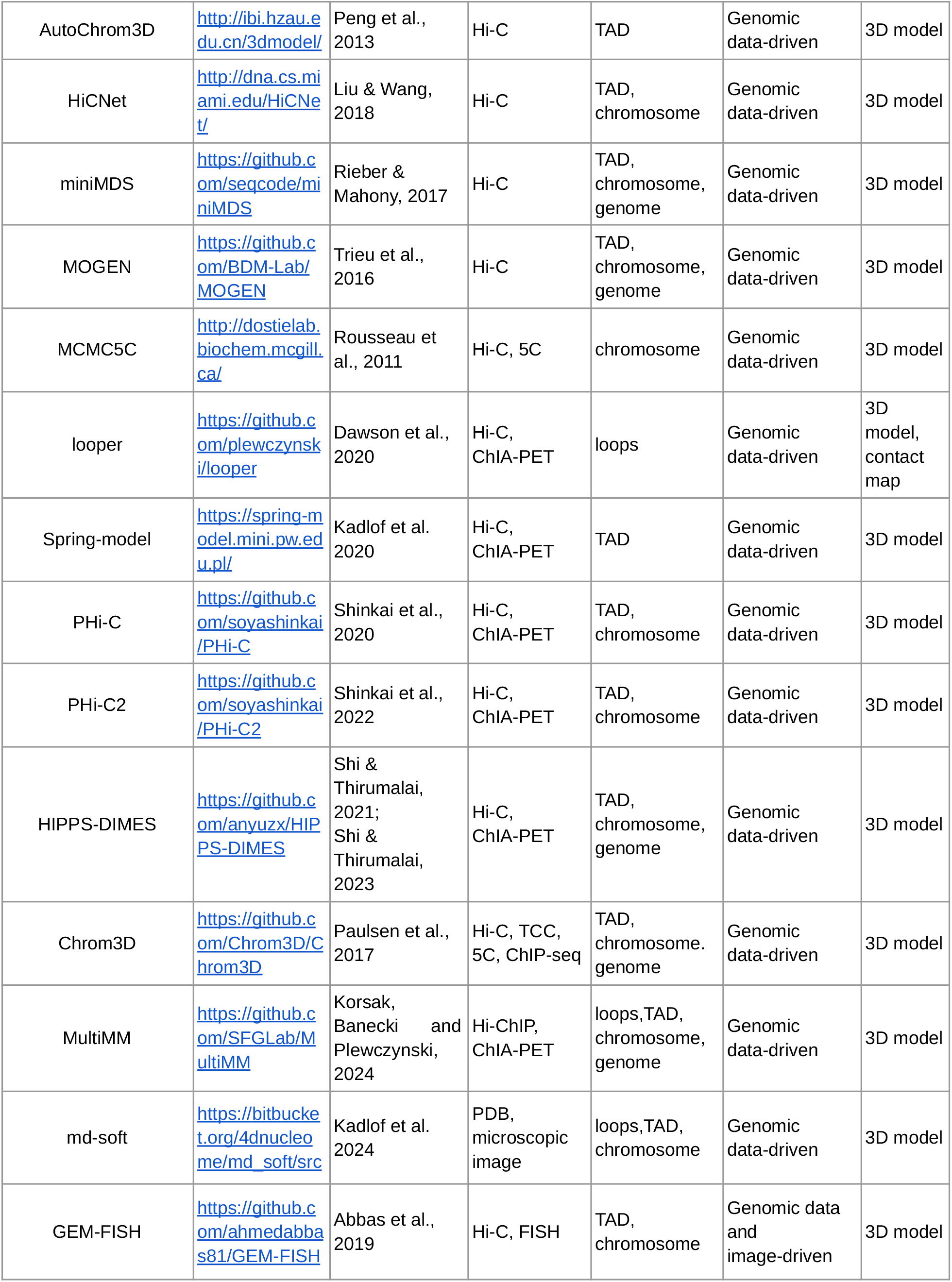
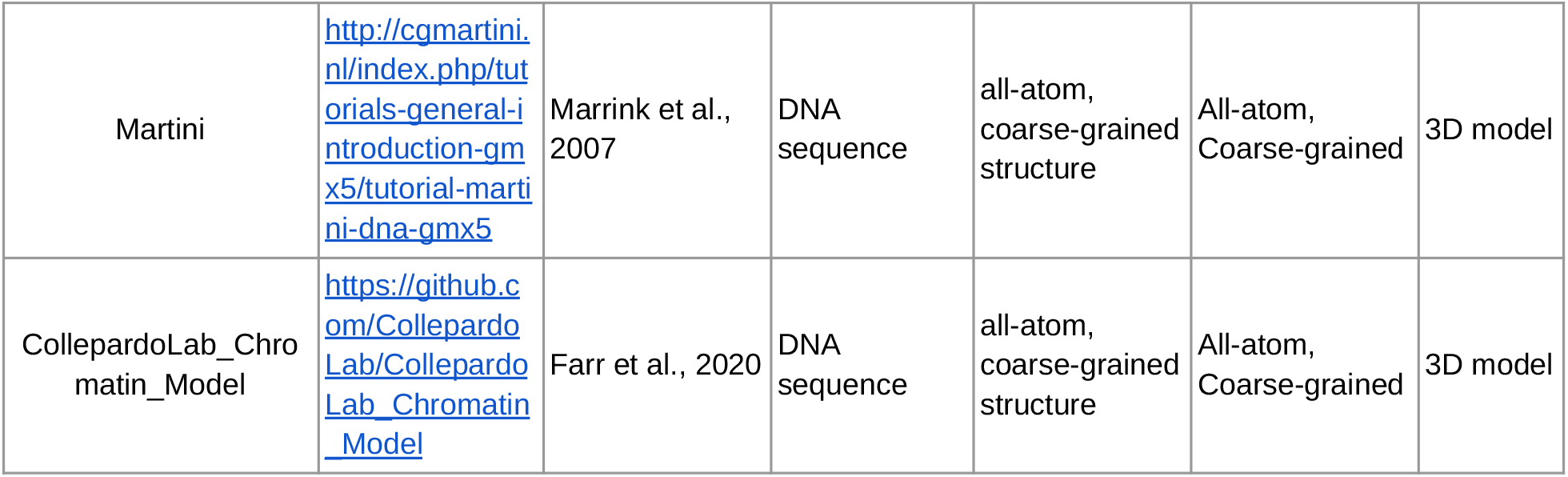
Overview of software for chromatin structure modeling. Software for chromatin modeling can be categorized into different categories based on input data, modeling scale, modeling strategy, or output format.

### 4D Nucleome Hackathon 2024

Hackathons are popular in life sciences, especially in the field of genomics, because they offer an amazing opportunity to foster international multi-disciplinary collaboration and to quickly advance projects based on the principle of open innovation (Walker et al., 2022; Deb et al., 2024). The advantage of doing software comparison in a hackathon setting is that one can collect diverse, unbiased, yet expert views on the software. Following this emerging idea, we participated in the 4D Nucleome Hackathon 2024, organized by the 4D Nucleome Consortium, that took place on March 18-21, 2024 at The University of Washington in Seattle, USA (event website: https://hack4dnucleome.github.io/). One of the 4D Nucleome Consortium aims is to study the structure and function of the human genome through predictive models of chromatin (Dekker et al., 2017; Dekker et al., 2023), however, currently, criteria for comparison and validation of such models are lacking, even though there have been initiatives undertaken to benchmark computational methods for chromatin modeling (Belokopytova & Fishman, 2021; Tao et al., 2021; International Nucleome Consortium, 2022). The hackathon offered an opportunity to define criteria for chromatin model comparison and validation, which, in theory, appears straightforward, however, it is problematic and complicated in practice. The objective of our hackathon project pertained to the investigation of diverse methodologies employed in the three-dimensional (3D) modeling of chromatin, each corresponding to divergent methodologies for the modeling of chromatin structure, with a concurrent comparison and examination of their usability across varied projects in life sciences. It is acknowledged that this undertaking was fraught with primary challenges, notably the dearth of experimental modalities required for method validation, compounded by the elevated computational intricacy stemming from the various scales and biophysical phenomena intricately intertwined with chromatin architecture. Indeed, achieving a complete theoretical comprehension of these biological processes presents an inherently formidable task.

Here we present a bioinformatic workflow to compare chromatin models and validate them against experimental data, which constitutes a way to progress in the modeling of the human genome. During the hackathon, due to a limited time frame and resources, we focused our efforts on five distinct software packages (DIMES, MultiMM, MiChroM, LoopSage, and PHi-C2), which allowed us to demonstrate the underlying challenges of chromatin structure modeling.

## Results

### Challenge of benchmarking

The main objective of the project was to address the challenges of chromatin model vs. model comparison, as well as the validation of chromatin models using experimental data (Hi-C (Lieberman-Aiden et al., 2009), ChIA-PET (Li et al., 2010) and SPRITE (Quinodoz et al., 2022)). Our hackathon experience highlighted several key challenges associated with benchmarking a large number of chromatin models. These challenges arise from multiple factors: (1) bioinformatic software is frequently developed in academic settings and often lacks long-term support, resulting in many outdated models with insufficient support and poor documentation (Anakella et al., 2017); (2) various models are designed to address different aspects of chromatin biophysics, focusing on diverse biophysical problems or scales, complicating direct comparisons; (3) the complexity of chromatin folding research necessitates expertise in biology, bioinformatics, and physics, which hinders the development of simple and user-friendly models. Despite these challenges, we successfully ran several modeling methods and developed a workflow for their benchmarking (Fig. 3).

**Fig. 3.**
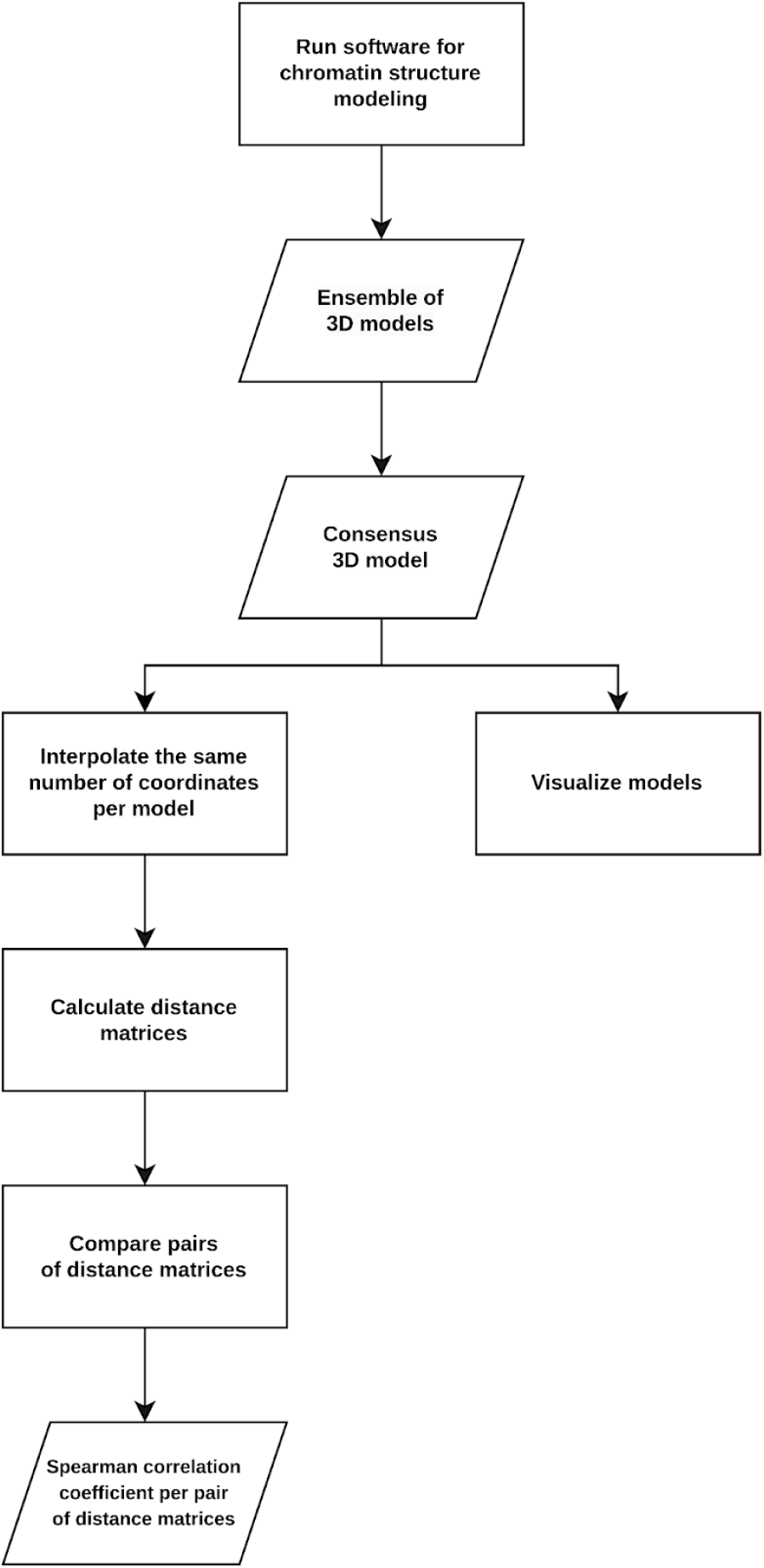
Workflow for benchmarking of chromatin structural models. The workflow developed at the 4D Nucleome Hackathon 2024 consists of multiple steps that include obtaining, visualizing, processing and comparing models of chromatin structure.

We began the project by executing each software listed in Table 1 consecutively and independently. If we encountered significant difficulties with a particular software, we opted to move on to the next one, bearing in mind the limited timeframe of the hackathon (4 days). The chromatin modeling software produced 3D models, which we visualized to examine the differences between them. Then we pursued the idea of converting these models into 2D contact matrices, which allowed us to calculate correlation coefficients for comparison, thus assigning a numerical value to each pair of matrices which represented the hypothesized differences between the models. We designed an easily programmable way to convert the models into matrices, and once we collected the matrices, we chose the Spearman correlation coefficient for matrix comparison. We discussed alternative metrics for comparison between two three-dimensional models (e.g., Pearson correlation coefficient or root mean squared deviation (RMSD)), however we decided to focus on Spearman correlation because of its simplicity, and the fact that it is easy to interpret, but also due to a limited time during the hackathon.

### Challenge of model vs. model comparison

During the 4-day hackathon, we followed our workflow to generate and compare chromatin models. We initially hypothesized that the models for the same genomic region would be consistent in shapes and sizes, therefore we chose a region of approximately 1 Mb that was of an appropriate length to model both short- and long-distance interactions. We selected two experimental data types Hi-C and ChIA-PET that provide information about genome folding in various cell types. Both of them come in the form of 2D contact matrices that we used as input for five distinct, user-friendly software packages: DIMES, MultiMM, MiChroM, LoopSage, and PHi-C2. We focused on the experimental data (Hi-C and ChIA-PET) for the Tier 1 *GM12878* human cell line and selected a chunk of it that corresponded to a topologically associated domain of approximately 1 Mb (chr1:178.421.513–179.491.193). We obtained output models (in XYZ, PDB, or CIF format) from those five software packages that generated either one model or an ensemble of models. We realized that the output models were of different resolutions denoting that they could not be directly compared. For that reason, we first interpolated each model to the same number of coordinates by finding an approximate basis spline representation of it, and obtained a uniform resolution across all of them. We settled for a final resolution of 214 beads for each model, equivalent to approximately 5000 base pairs per bead, however, for the lack of time we did not assess how much information was lost during the interpolation, nor did we pursue the idea of testing other values for resolution. Nonetheless, with those standardized models, we were able to move forward with the project and create distance matrices of consistent shapes for all of them. We visualized the output models of the genomic region of interest (chr1:178.421.513–179.491.193) generated using Hi-C data (Fig. 4), as well as ChIA-PET data (Fig. 5), however, the models displayed inconsistencies in both shapes and sizes. Therefore, we considered correlation coefficients, but not the model size, as the comparison metric. Despite using the same input data for all software, the reason for the disparities might be due to the differences in modeling methodologies and assumptions used in the software. As previously mentioned, algorithms are designed to address specific aspects of chromatin biophysics and focus on different biological scales, complicating model comparisons. Moreover, we acknowledge that we used default parameters for all software packages, therefore leaving open the possibility that the parameters could have influenced the modeling process and results.

**Fig. 4.**
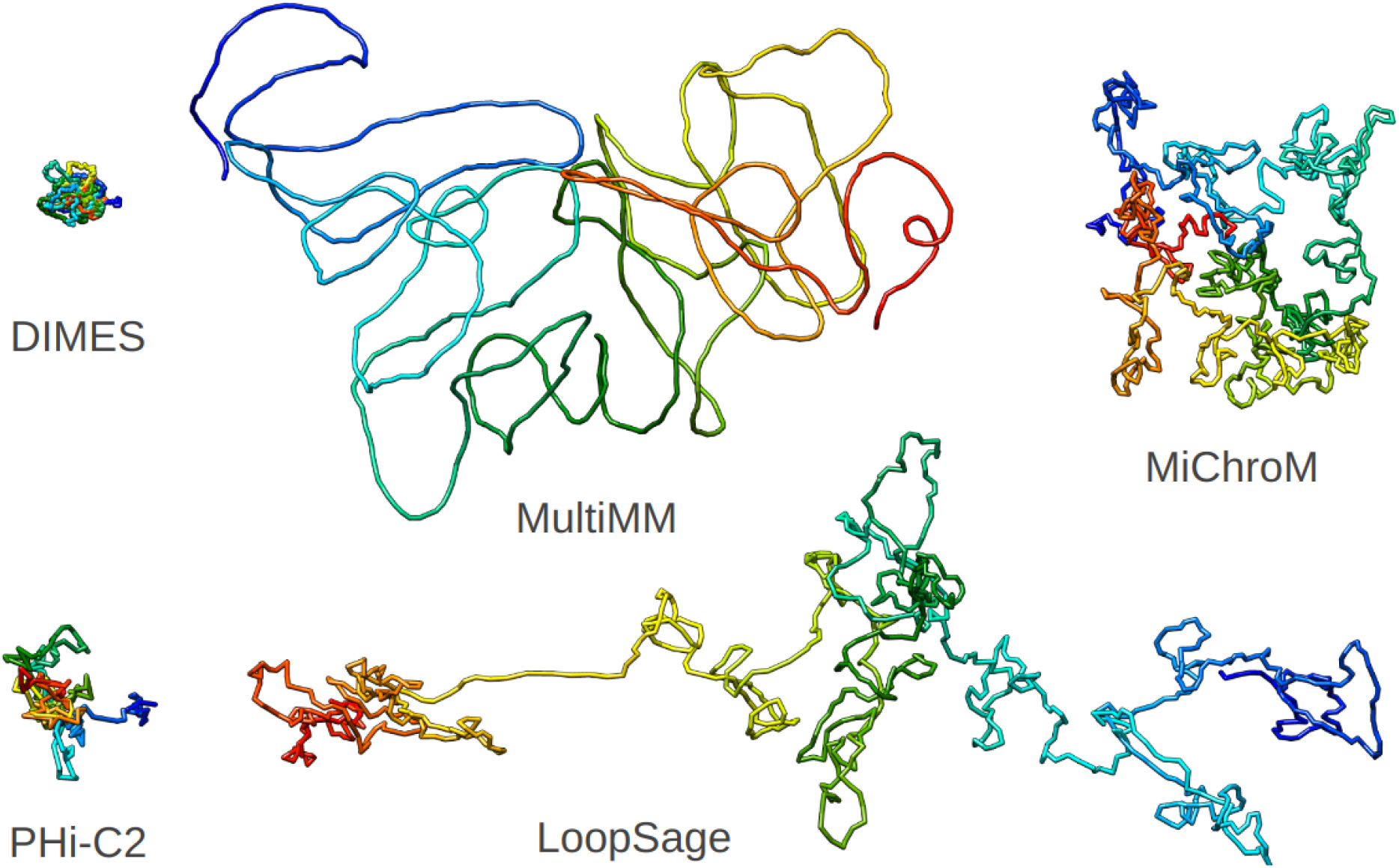
Chromatin models showing the same topologically associated domain. Models presenting the same genomic region (chr1:178.421.513–179.491.193) obtained from five software packages (DIMES, MultiMM, MiChroM, LoopSage, and PHi-C2) based on Hi-C data.

**Fig. 5.**
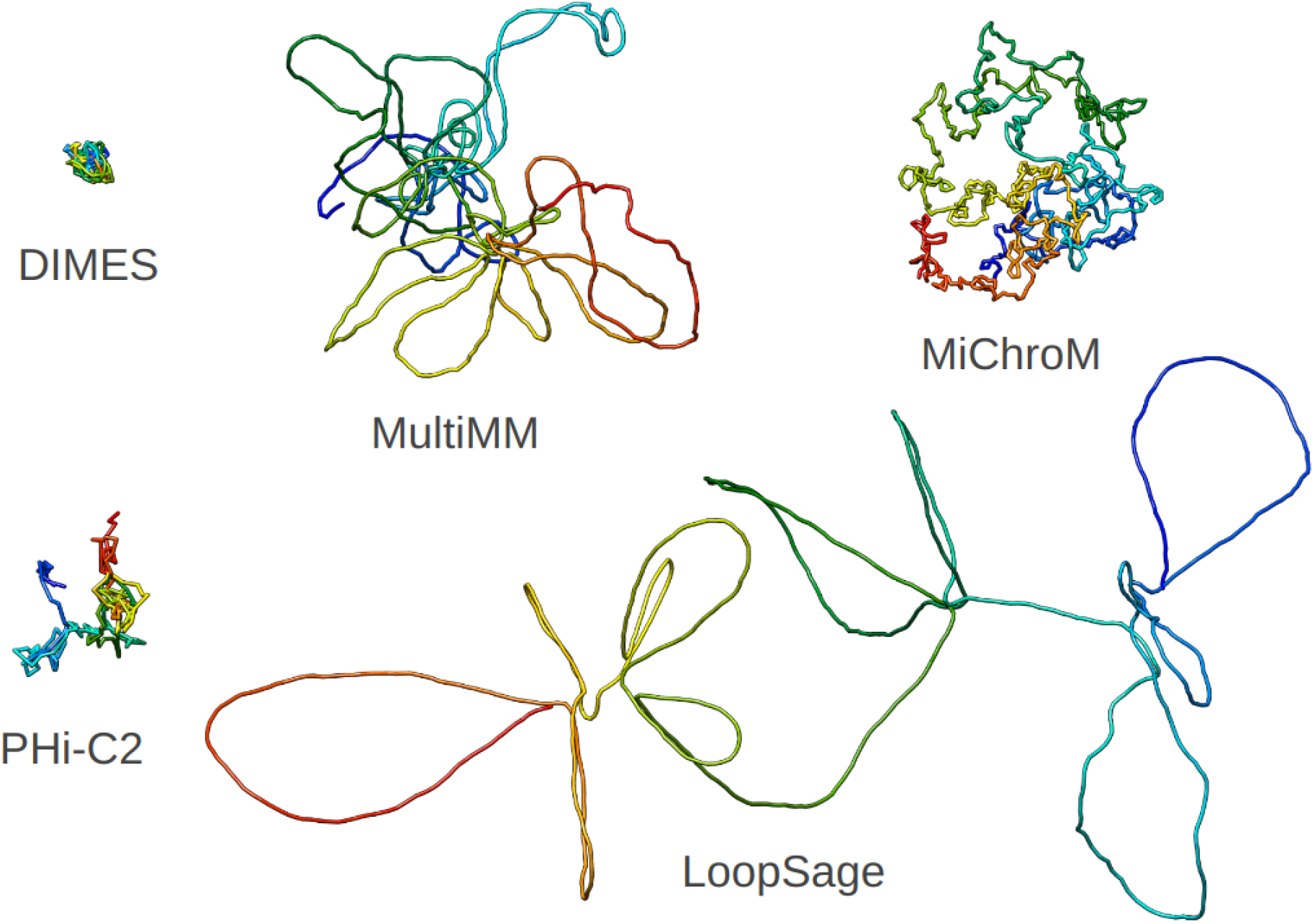
Chromatin models showing the same topologically associated domain. Models presenting the same genomic region (chr1:178.421.513–179.491.193) obtained from five software packages (DIMES, MultiMM, MiChroM, LoopSage, and PHi-C2) based on ChIA-PET data.

We hypothesized that if, in addition to Hi-C data, we used ChIA-PET data, which provides different insights about genome folding compared to Hi-C data, we would obtain different models. This assumption comes from the complementary, but not indistinguishable, nature of both experimental methods. Both can be represented as contact matrices, however, the elements of Hi-C matrices represent contact probabilities between genomic regions, whereas ChIA-PET matrices represent genomic interactions mediated by proteins involved in genome folding. Therefore, we used the same software once more, however we provided ChIA-PET data as input. The output model visualizations demonstrate that first, all ChIA-PET models differ from each other in shapes, but similarly using ChIA-PET instead of Hi-C does not yield identical structures (Fig. 5). These findings highlight the challenges of chromatin structure modeling and emphasize the need for robust methods to compare models. To address this, we proposed converting the 3D models into distance matrices, allowing us to quantify the differences between them through matrix comparisons.

In the next step, for all models, we computed 2D distance matrices, the elements of which corresponded to average pairwise Euclidean distances between the beads. Since certain software produces ensembles of models, then the elements of the 2D matrix are averaged over all models in the ensemble. Otherwise, only one structure was represented as the 2D matrix. We visualized the matrices as heatmaps, then we set the heatmaps side by side, and we noticed that the differences observed in visualizations are reflected in heatmaps as well (Fig. 6).

**Fig. 6.**
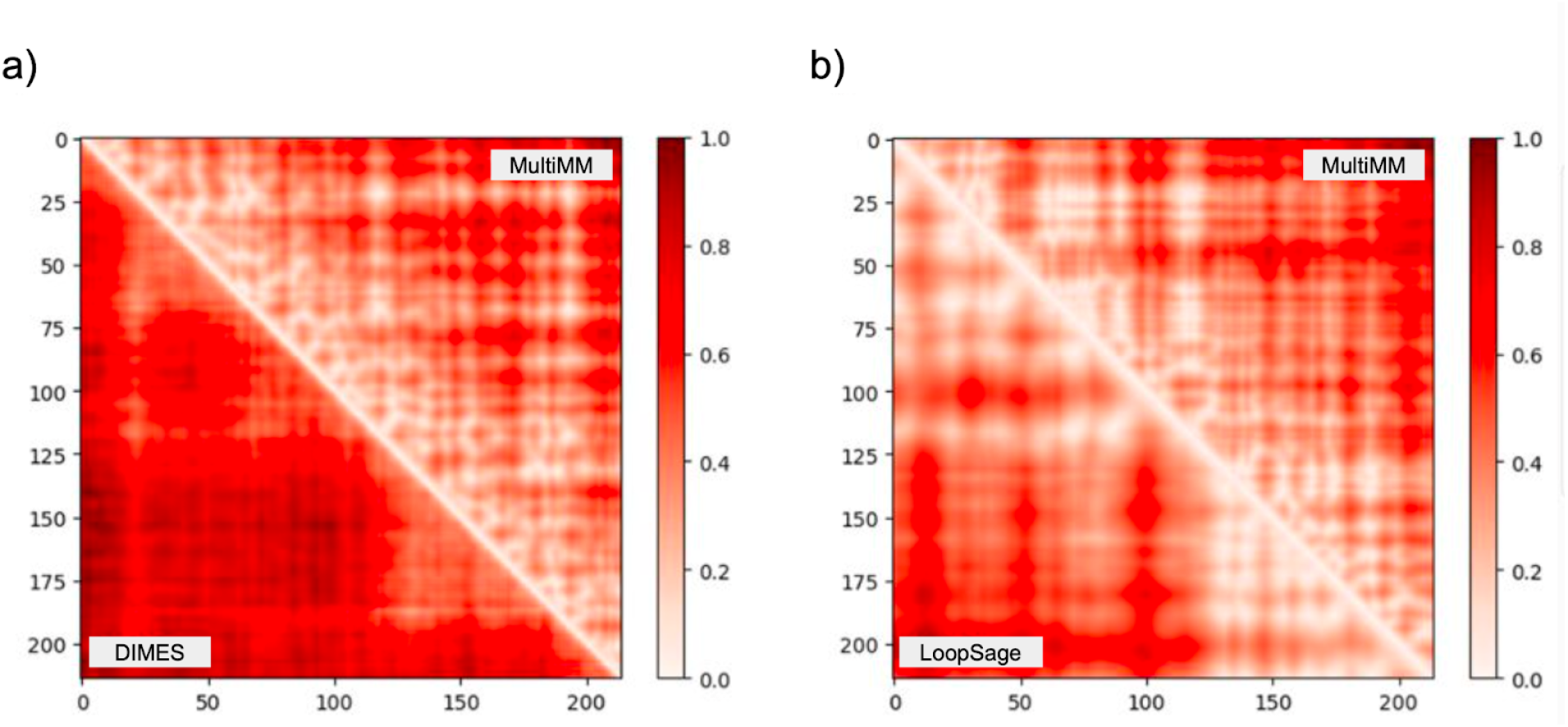
Comparisons of heatmaps generated from the models based on Hi-C data: a) from DIMES and MultiMM, b) from LoopSage and MultiMM. All heatmaps correspond to the interpolated models (number of beads = 214) that were generated for the same region of interest (chr1:178.421.513–179.491.193).

Finally, the matrices allowed us to calculate a Spearman correlation coefficient (ρ) (that takes a value between -1 and 1) for each pair of matrices to assess similarities between the corresponding models, while the higher the value of the Spearman correlation coefficient, the more similarity there was between the models in terms of the spatial distribution of beads. We recognized and accepted the possibility that other similar metrics (e.g., Pearson correlation coefficient) or combinations thereof might be more appropriate for matrix comparison, however, we selected the most straightforward and intuitive approach for the purpose of the hackathon. We expected to observe a rather significant heterogeneity in the coefficients since the models differed notably in shapes, and as anticipated, the coefficients reflected the discrepancies between the models. In terms of the models generated with Hi-C data (Table 2), the highest correlation was observed between the models generated with MultiMM and DIMES (ρ = 0.725), while the lowest correlation was observed between the models from PHi-C2 and MiChroM (ρ = 0.373). As regards the models generated from ChIA-PET data (Table 3), the highest correlation was found between the models generated with LoopSage and MultiMM (ρ = 0.800), while the lowest correlation was shown to be between the models from MiChroM and DIMES (ρ = 0.248). Overall, by using Spearman correlation coefficients, we were able to demonstrate a great heterogeneity in chromatin models produced by the tested software packages that were also observed in model visualizations.

**Table 2.** Comparison of chromatin models generated based on Hi-C data. The table contains Spearman correlation coefficients calculated between pairs of distance matrices generated based on the models from five software packages (DIMES, MultiMM, MiChroM, LoopSage, and PHi-C2).

| DIMES |  |  |  |  |  |
| --- | --- | --- | --- | --- | --- |
| MultiMM | 0.725 |  |  |  |  |
| MiChroM | 0.406 | 0.382 |  |  |  |
| LoopSage | 0.643 | 0.626 | 0.448 |  |  |
| PHi-C2 | 0.593 | 0.574 | 0.373 | 0.610 |  |
|  | DIMES | MultiMM | MiChroM | LoopSage | PHi-C2 |

**Table 3.** Comparison of chromatin models generated based on ChIA-PET data. The table contains Spearman correlation coefficients calculated between pairs of distance matrices generated based on the models from five software packages (DIMES, MultiMM, MiChroM, LoopSage, and PHi-C2).

| DIMES |  |  |  |  |  |
| --- | --- | --- | --- | --- | --- |
| MultiMM | 0.275 |  |  |  |  |
| MiChroM | 0.248 | 0.413 |  |  |  |
| LoopSage | 0.341 | 0.800 | 0.418 |  |  |
| PHi-C2 | 0.365 | 0.776 | 0.413 | 0.751 |  |
|  | DIMES | MultiMM | MiChroM | LoopSage | PHi-C2 |

### Challenge of model validation

The second challenge we decided to address during the hackathon was the validation and interpretation of chromatin models. Such models aim to bridge theory and experiment, therefore it is crucial to understand how experimental data underlies distances between genomic regions in the model and how close the model is to the real chromatin structure. This would advance the design of future experiments that aim to study the impact of the genome structure, i.e., proximity of various genomic regions (e.g., genes, promoters and enhancers) on gene expression and other cellular processes. During the hackathon, it was not easy to formulate the exact hypothesis and define criteria for model validation. Firstly, chromatin models represent spatial distances between genomic regions, while experimental data can show contact probabilities (Hi-C), genomic interactions mediated by proteins involved in genome folding (ChIA-PET) or high-order genomic interactions (SPRITE). Those experimental methods complement each other, however they provide different biological information. Furthermore, the difficulty of interpreting experimental data itself further impedes the challenge of model validation. Finally, there are currently no standard criteria or metrics to conduct such validation.

Our project demonstrates that model validation is indeed a difficult task, even with expertise in both software and experimental data analysis. Here, we present our approach for model validation, in which we convert models into distance matrices, and then calculate Spearman correlation coefficients (ρ) between them to quantify model similarities. During the hackathon, we examined how models generated using Hi-C and ChIA-PET data correlate with three different biological data sets: Hi-C, ChIA-PET and SPRITE. To do this, we used the same chromatin models that we previously generated using Hi-C and ChIA-PET data for model comparison. We hypothesized that the distance matrices generated from Hi-C models would correlate more strongly with Hi-C data, while the matrices from ChIA-PET models would show a higher correlation with ChIA-PET data. To test our hypothesis, we calculated Spearman correlation coefficients between model matrices and the inverse of experimental contact frequency matrices. Our analysis revealed a significant heterogeneity in Spearman correlation coefficients across both the software packages, as well as the biological data sets (see Supplementary Information). For instance, we observed the highest correlation between DIMES and Hi-C (ρ = 0.731) and the lowest correlation between MiChroM and SPRITE (ρ = 0.185). For models generated using Hi-C data, we observed a consistent pattern: they tended to correlate more strongly with Hi-C data compared to ChIA-PET or SPRITE data.

Moreover, the Hi-C models correlated well with Hi-C data, but interestingly, the ChIA-PET models also showed a stronger correlation with Hi-C data rather than ChIA-PET data. These results provide insights into the challenge of chromatin model validation and demonstrate that the Spearman correlation coefficient can constitute a metric that represents how models are correlated with experimental data. However, this validation methodology requires further investigation, for instance, to examine how other similarity metrics and combinations thereof would fit into our approach. Moreover, it would be worthwhile to examine how both machine learning and biophysics-based software, which differ in approaches, assumptions, scales and input data (e.g., imaging data) correlate with experimental data.

## Discussion

Prior to the hackathon, we identified that there is a lack of objective metrics to compare and validate 3D models of chromatin structure. It has been previously discussed that software performance, usability and interpretability are key aspects for studying genome folding (Di Stefano et al., 2021, Belokopytova & Fishman, 2021; Tao et al., 2021; Liu et al., 2023), therefore we set an aim for the 4D Nucleome Hackathon 2024 to develop a bioinformatics workflow for model comparison and validation. We started with a thorough literature review on available software packages for chromatin structure modeling that use various experimental data types, either Hi-C, ChIA-PET, ChIP-seq, imaging data or combination thereof. In the end, we gathered a number of software packages that were later classified by the scale of modeling, starting from the smallest scale - loops and TADs, through chromosomes to the whole genome. Throughout the process we got an overview of the variety of approaches that have been undertaken to predict the structure of chromatin, which can be either physics- or machine learning-based. There is a great number of approaches in software for chromatin structure modeling, as well as several shortcomings of those methods: (1) bioinformatic software frequently lacks long-term support and informative documentation (Anakella et al., 2017); (2) software is usually designed to address specific aspects of chromatin biophysics, focusing on diverse biophysical problems or scales, which complicates software comparisons; (3) the complexity of chromatin folding necessitates expertise in biology, bioinformatics, and physics; (4) software for chromatin structure modeling requires objective metrics to quantify its efficiency.

**The first challenge** of our project was that a great number of software is not open-source, nor runnable without detailed technical knowledge and the field lacked a general and formal standardization. Our results indicate that it is indeed a difficult problem, and therefore we emphasize an unidentified need for software accessibility and reproducibility, that would lower the entry barrier for young researchers to enter the field, thus enabling a quicker implementation of novel innovative ideas. In addition, there are no common guidelines for software development. We therefore hypothesize that standardization and guidelines for software development, which are currently challenging to define, would have a positive long-term impact on the community.

Moreover, **the second challenge** in the field of structural genomics is the lack of objective criteria for model comparison and validation. To address this challenge, we present a modular and scalable workflow for processing and comparing chromatin models that includes a conversion of models into distance matrices and calculation of Spearman correlation coefficients between pairs of matrices that represent similarities between them. As a proof-of-principle, during the hackathon, we compared models of one genomic region (a topological associated domain of 1 Mb), obtained from five distinct software packages. We identified a big heterogeneity in the output models, which might be due to the variety of approaches and assumptions in the software, however we acknowledge that this undertaking might be fraught with primary challenges, which were due to a limited time frame and resources. Nevertheless, we believe that our workflow provides a future reference for other initiatives that might be undertaken to develop criteria for chromatin model comparison and validation. For that purpose, we made our workflow publicly available on GitHub (https://github.com/SFGLab/Polymer_model_benchmark) and to ensure reproducibility, we provide scripts and virtual environment files to run it on any Linux/GNU-based computing system.

Looking forward, it would be worthwhile to do a comprehensive study of all software for chromatin modeling, and especially to include 3D genomic methods incorporating artificial intelligence and single-cell technologies, therefore we plan to extend our joint effort to focus on those methodologies as well. For that reason, it would be crucial to examine how those novel technologies can advance the modeling itself, as well as the downstream analyses and model interpretation. Another potential avenue for improvement of chromatin modeling methods might be sought in the integration of 3D genomics data with multi-omic next-generation sequencing data to study the impact of genomic variation on the genome structure and function. To conclude, here we emphasize that chromatin modeling is crucial for biological and biomedical research, as well as identify and discuss the challenges that impede usability, reproducibility, and interpretability of the software for chromatin modeling.

## Methods

### Software

During the hackathon we used the following five software packages that incorporate different underlying methodologies based on various biophysical principles:

- LoopSage (Korsak & Plewczynski 2024; https://github.com/SFGLab/LoopSage),
- MiChroM (Di Pierro et al., 2016; https://open-michrom.readthedocs.io/en/latest/OpenMiChroM.html),
- DIMES (Shi & Thirumalai, 2023; https://github.com/anyuzx/HIPPS-DIMES),
- PHi-C2 (Shinkai et al., 2020; https://github.com/soyashinkai/PHi-C2),
- MultiMM (Korsak, Banecki and Plewczynski, 2024; https://github.com/SFGLab/MultiMM).

Our workflow was implemented in Python (v3.12.2) using the NumPy (v1.26.4) and SciPy (v1.12.0) libraries.

### Data

Comparing various modeling techniques poses inherent challenges, necessitating the proposition of a methodologically straightforward approach for comparison. Initially, our focus was to evaluate the performance of these models within small-scale topologically associated domain (TAD) regions and to assess their congruence with experimental data. In order to model the whole genome, a huge amount of computational resources, as well as time would be required. Due to the lack of those resources during the 4-day hackathon, a short genomic region of interest (chr1:178.421.513–179.491.193) for the Tier 1 cell line *GM12878* was chosen to generate the models. It is approximately 1 Mb long, and it represents a topologically-associated domain (TAD). We downloaded public data from the 4DNucleome Data Portal (https://data.4dnucleome.org/) (Dekker et al., 2017; Reiff et al., 2022) and ENCODE (https://www.encodeproject.org/) (The ENCODE Project Consortium 2012 ; Luo et al., 2020 ; Meenakshi et al., 2023). Chromatin models were generated based on the following *in situ* Hi-C data from the 4DN Data Portal: 4DNES4AABNEZ, 4DNESNMAAN97 and ENCODE: ENCSR968KAY, as well as ChIA-PET from ENCODE: ENCSR184YZV, ENCSR764VXA. For model validation, we used SPRITE data from 4DN Data Portal: 4DNESI1U7ZW9.

### Code and data availability

To ensure that these results are reproducible, all scripts for model comparison and validation have been made publicly available on GitHub: https://github.com/SFGLab/Polymer_model_benchmark

## Acknowledgements

We thank The 4D Nucleome Consortium for organizing and sponsoring the 4D Nucleome Hackathon 2024, and The University of Washington for hosting the event. DP, JK, MK, SK, KB research was funded by Warsaw University of Technology within the Excellence Initiative: Research University (IDUB) programme, their work has been co-supported by Polish National Science Centre (2020/37/B/NZ2/03757) and the National Institute of Health USA 4DNucleome grant 1U54DK107967-01 “*Nucleome Positioning System for Spatiotemporal Genome Organization and Regulation*”. Part of the high-performance computations were performed thanks to the Laboratory of Bioinformatics and Computational Genomics, Faculty of Mathematics and Information Science, Warsaw University of Technology using Artificial Intelligence HPC platform financed by Polish Ministry of Science and Higher Education (decision no. 7054/IA/SP/2020 of 2020-08-28).

