## Supplementary Table for "The challenge of chromatin model comparison and validation - a project from the first international 4D Nucleome Hackathon"

### Supplementary Information

**Table 1. Comparison of chromatin models generated based on Hi-C with experimental data.** The table contains Spearman correlation coefficients calculated between the distance matrices generated based on the models from with five software packages (DIMES, MultiMM, MiChroM, LoopSage, and PHi-C2) with three experimental data types (Hi-C, ChIA-PET, and SPRITE).

| | **DIMES** | **MultiMM** | **MiChroM** | **LoopSage** | **PHi-C2** |
|---|---|---|---|---|---|
| **Hi-C** | 0.731 | 0.534 | 0.340 | 0.501 | 0.496 |
| **ChIA-PET** | 0.502 | 0.411 | 0.293 | 0.388 | 0.415 |
| **SPRITE** | 0.393 | 0.308 | 0.185 | 0.308 | 0.211 |

**Table 2. Comparison of chromatin models generated based on ChIA-PET with experimental data.** The table contains Spearman correlation coefficients calculated between the distance matrices generated based on the models from with five software packages (DIMES, MultiMM, MiChroM, LoopSage, and PHi-C2) with three experimental data types (Hi-C, ChIA-PET, and SPRITE).

| | **DIMES** | **MultiMM** | **MiChroM** | **LoopSage** | **PHi-C2** |
|---|---|---|---|---|---|
| **Hi-C** | 0.332 | 0.554 | 0.342 | 0.576 | 0.608 |
| **ChIA-PET** | 0.217 | 0.437 | 0.281 | 0.445 | 0.504 |
| **SPRITE** | 0.16 | 0.33 | 0.147 | 0.339 | 0.328 |
